# The integrity and assay performance of tissue mitochondrial DNA is considerably affected by choice of isolation method

**DOI:** 10.1101/2021.02.10.430587

**Authors:** Bruno Marçal Repolês, Choco Michael Gorospe, Phong Tran, Anna Karin Nilsson, Paulina H. Wanrooij

## Abstract

The integrity of mitochondrial DNA (mtDNA) isolated from solid tissues is critical for analyses such as long-range PCR, but is typically assessed under conditions that fail to provide information on the individual mtDNA strands. Using denaturing gel electrophoresis, we show that commonly-used isolation procedures generate mtDNA containing several single-strand breaks per strand. Through systematic comparison of DNA isolation methods, we identify a procedure yielding the highest integrity of mtDNA that we demonstrate displays improved performance in downstream assays. Our results highlight the importance of isolation method choice, and serve as a resource to researchers requiring high-quality mtDNA from solid tissues.

## Introduction

Mammalian mitochondrial DNA (mtDNA) is a double-stranded multicopy genome of approximately 16.5 kbp. Because it contains the genetic information encoding for core subunits of the mitochondrial respiratory chain, the integrity of mtDNA is critical to the normal function of the majority of mammalian cells. Alterations in mtDNA, such as mutations or large-scale deletions, can cause mitochondrial disease and are also implicated in a wide range of other pathologies as well as in the normal process of ageing (Szczepanowska and Trifunovic, 2017; Taylor and Turnbull, 2005; Wallace, 2012). For this reason, the accumulation, metabolic consequences and potential repair mechanisms of mutations or various types of damage in mtDNA are subject to intense research.

Many pertinent research efforts in the field rely on meticulous analysis of mtDNA isolated from solid mammalian tissues. Depending on the requirements dictated by downstream analysis methods, most researchers opt for a DNA isolation method that belongs to one of two main categories: isolation of mtDNA from purified or enriched mitochondria, or, if the planned analysis can distinguish mtDNA from a pool of other nucleic acids, the isolation of total genomic DNA (*i.e*. nuclear DNA and mtDNA). Both approaches have their benefits and disadvantages. Isolation of mtDNA from purified mitochondria yields highly-enriched mtDNA, but is more labor intensive, time consuming, and requires larger amounts of tissue as starting material. Moreover, the tissue homogenization step may require adjustment depending on the type of tissue used. In comparison, isolation of total DNA is faster and more straight forward as it requires little optimization to the type of tissue. It produces sufficient DNA yields from a smaller mass of tissue and is therefore often the method of choice. Many variants of the procedure exist, most of which are easily scaled up for the simultaneous handling of multiple parallel samples. Tissue lysis is often achieved with the help of a detergent like SDS in conjunction with proteinase K in an EDTA-containing buffer, conditions that inactivate most cellular endonucleases (Lang and Burger, 2007).

Comparisons of various mtDNA isolation techniques typically only analyze the mtDNA under neutral conditions, which does not yield information on the integrity of the individual strands and thereby prevents comparison of the frequency of single-stranded DNA (ssDNA) breaks. We show that the choice of isolation method has a considerable impact on the integrity of the mtDNA isolated from solid mammalian tissues, and that it further affects the performance of downstream assays. A systematic comparison of four different isolation procedures with regard to the integrity, yield and enrichment of the resulting mtDNA identifies a preferred isolation method for applications that require tissue mtDNA of high integrity.

## Materials and Methods

### Animal handling and isolation of tissues

Experiments were carried out using surplus wildtype C57BL/6 animals from other studies. All mice were maintained at the animal facility at Umeå University under pathogen-free conditions. Mice were housed in an environment with a 12-hour dark/light cycle and *ad libitum* access to food and water. Animal handling and experimental procedure were approved by the ethical committee at Umeå University, complying with the rules and regulations of the Swedish Animal Welfare Agency and with the European Communities Council Directive of 22 September 2010 (2010/63/EU). All efforts were made to minimize animal suffering and to reduce the number of animals used.

Mice were euthanized by CO_2_ inhalation at the age of 13-16 weeks (‘adult’) or 23–29 months (‘aged’). Livers, hearts, brains and hind leg muscles (gastrocnemius [GAS], tibialis anterior [TA], and mixed thigh muscles) were placed in Eppendorf tubes, quickly frozen in liquid N_2_ (unless indicated otherwise) and kept at −80°C. As far as possible, samples from an equal number of male and female animals were used in each experiment.

### DNA isolation and treatments

In the standard isolation method (Sambrook and Russell, 2001), ca 100 mg of individual mouse tissue was cut into small pieces, incubated at 55°C overnight in SNET lysis buffer (20 mM Tris pH 8.0, 5 mM EDTA pH 8.0, 400 mM NaCl, 1% SDS) containing 400 μg/ml proteinase K (#P6556, Sigma), and total DNA (*i.e.* nuclear DNA and mtDNA) was isolated by phenol-chloroform extraction and isopropanol precipitation according to standard protocols (Green and Sambrook, 2017).

In the pulverization method, ca 100 mg of frozen tissue was ground to a powder in a mortar containing liquid N_2_. The pulverized tissue was incubated in SNET buffer and proteinase K at 55°C for 3 h, followed by phenol-chloroform extraction and isopropanol precipitation as above. Samples were treated with 1.2 μg/μl PureLink RNase A (Invitrogen, #12091-021) at 37°C for 30 min and precipitated again with isopropanol.

In the kit-purification method, total DNA was extracted using the NucleoSpin Tissue kit from Macherey-Nagel (#740952) according to the manufacturer’s instructions. Briefly, 100 mg of tissue was cut into small pieces and incubated at 56°C for 3 h in Buffer T1 with Proteinase K, followed by a 5-minute incubation with RNase A at room temperature. Samples were lysed with 1 volume of Buffer B3 and incubated at 70°C for 10 min. After addition of 1 volume ethanol, the samples were applied to the NucleoSpin Tissue columns and centrifuged for 1 min at 11,000 x *g*. Silica membranes were washed twice with 500 μl Buffer BW with centrifugation for 1 min at 11, 000 x *g*, centrifuged once more to dry the silica membrane and remove residual ethanol, and DNA eluted with 30 μl TE buffer (10 mM Tris pH 8.0; 1 mM EDTA).

Isolation of mtDNA from enriched mitochondria was modified from (Ekstrand et al., 2004; Frezza et al., 2007; Yasukawa et al., 2005) using ca 200-300 mg of finely-chopped tissue. The liver samples were washed thrice in homogenization buffer (HB; 225 mM mannitol, 75 mM sucrose, 10 mM Hepes-KOH pH 7.8, 10 mM EDTA), while skeletal muscle samples were washed in ice-cold PBS containing 10 mM EDTA, incubated with 0.05% trypsin (w/v) in PBS with 10 mM EDTA for 90 min on ice, centrifuged (200 × *g*, 5 min, 4°C), and the pellet resuspended in HB. Tissues were homogenized by 10 strokes in a hand-held glass/Teflon Potter Elvehjem homogenizer. Nuclei and cell debris were pelleted at 800 × *g* for 10 min at 4°C and the supernatant was further centrifuged at 12,000 × *g* for 10 min at 4°C to pellet mitochondria. The mitochondrial pellet was resuspended in HB buffer containing 100 ng/μl proteinase K and incubated for 15 min on ice. Mitochondria were pelleted (12,000 × *g*, 10 min, 4°C), resuspended in lysis buffer (75 mM NaCl, 50 mM EDTA, 1% SDS, 20 mM Hepes pH 7.8) containing 100 ng/μl proteinase K, and incubated on ice for 30 min. MtDNA was purified by phenol–chloroform extraction followed by ethanol precipitation.

All DNA preparations were resuspended in TE and allowed to dissolve overnight. In order to minimize DNA shearing, DNA samples were not vortexed at high speed, and cut or wide-bore tips were used for pipetting whenever feasible. DNA concentrations were determined by Nanodrop. Where indicated, DNA preparations were digested overnight with the SacI restriction endonuclease (Thermo Scientific, #ER1131), followed by ethanol precipitation and resuspension in TE buffer. T4 DNA ligase treatment (Thermo Scientific, #EL0011; at 3.3 U/μg of DNA) was carried out at 25°C for 6 h, followed by inactivation at 65°C for 10 min in the presence of 25 mM EDTA and 0.2% SDS.

### Agarose gel electrophoresis and Southern blotting

Pure mtDNA (500 ng/lane) and total DNA (2 μg/lane) samples were separated on 0.7% agarose gels under either neutral (1 × TAE buffer) or denaturing (30 mM NaOH and 1 mM EDTA) conditions at 1.2 V/cm for 16-18 h at 8°C; 5 ng of GeneRuler 1 kb DNA ladder (Thermo Scientific) was used as a standard for DNA length. The DNA was transferred to a nylon membrane using standard protocols (Sambrook and Russell, 2001) and probed sequentially with selected α-dCTP^32^-labelled dsDNA probes, stripping extensively with boiling 0.1% SDS between probes. The probes used were *COX1* (nt 5,328–6,872) for mouse mtDNA and 18S (nt 1,245–1,787 of the 18S rRNA on chromosome 6 [gi 374088232]) for nuclear DNA. Blots were exposed to BAS-MS imaging plates (Fujifilm), scanned on a Typhoon 9400 imager (Amersham Biosciences), and the signal was quantified by using ImageJ64 software. Length distributions of individual DNA samples were used to determine the apparent median length of DNA fragments as previously described (Wanrooij *et al.*, 2017). Unless otherwise noted, statistical comparisons were performed by using Welch’s unequal variances t-test (for comparisons between two groups) or one-way ANOVA (between three or more groups) in GraphPad Prism software.

### Analysis of mtDNA copy number (enrichment) by qPCR

MtDNA copy number was analyzed in duplicate by quantitative real-time PCR using 1 μl of 1/200-diluted DNA in a 20 μl reaction containing 0.2 μM forward and reverse primers and 10 μl of 2x SyGreen Mix (qPCRBIO #PB20.14-05) in a LightCycler 96 instrument (Roche). Primer pairs targeting cytochrome B (nt 14,682–14,771 of mtDNA; forward GCTTTCCACTTCA TCTTACCATTT, reverse TGTTGGGTTGTTTGATCCTG (Ahola-Erkkilä *et al.*, 2010)) and actin (nt 142,904,143–142,904,053 of chromosome 5 [NC_000071]; forward CCACCATGTACCCAGGC, reverse CACCCTCACCAAGCTAAGG) were used with the following qPCR program: 95°C 180 sec, 45 cycles of (95°C 10 sec, 57°C 10 sec, 72°C 20 sec with signal acquisition), melting curve (95°C 5 sec, 65°C 60 sec, heating to 97°C at 0.2°C/sec with continuous signal acquisition). Cq values determined by the LightCycler 96 software (Roche) were used to calculate the copy number of mtDNA relative to nuclear DNA using the Pfaffl method (Pfaffl, 2001) and plotted with GraphPad Prism. Statistical comparisons between multiple groups were performed by one-way ANOVA.

### Long-range quantitative PCR

Long-range PCR for an 13,481-bp fragment of mtDNA was performed by quantitative real-time PCR using 6.25 ng of DNA, 0.2 μM each of forward (2,478F: GTTTGTTCAACGATTAAAGTCCTACGTGATCTGAG) and reverse (15,958R: GCCTTGACGGCTATGTTGAT) primers, 200 μM dNTPs, 1x concentration of EvaGreen Dye (Biotium, #31019) and 0.63 U PrimeSTAR GXL polymerase (Takara, #R050A) in the corresponding PrimeSTAR GXL buffer. Reactions were performed in duplicate in a LightCycler 96 instrument (Roche) with the following program: 95°C 180 sec; 35 cycles of (95°C for 30 sec, 68°C for 10 min with signal acquisition), melting curve (95°C 5 sec, 65°C 60 sec, heating to 97°C at 0.2°C/sec with continuous signal acquisition). As internal control, a 145-bp fragment was amplified from each sample using the same reverse primer, an internal forward primer (15,814F: AAATGCGTTATCGCCCATAC), and the same PCR program with the exception of a 1-min amplification time. C_q_ values of both amplicons were used to compute the relative amplification of the long fragment relative to the short fragment, normalizing values to the average of the samples prepared using the standard isolation method. Statistical comparison of each group to the standard group was performed by one-way ANOVA.

## Results

### Tissue mtDNA isolated by standard methods contains single-stranded DNA breaks

We isolated total genomic DNA from the skeletal muscle (tibialis anterior [TA]) and the liver of adult (13-16-week-old) and aged (2-year-old) mice using a standard DNA isolation protocol (Sambrook and Russell, 2001) and carried out gel analysis under two different conditions: neutral conditions where the mtDNA remains in double-stranded form, and denaturing conditions that separate the two strands. For unambiguous interpretation of the results, the DNA separated under neutral conditions was first treated with SacI endonuclease to linearize the mtDNA. The mtDNA was then visualized by Southern blotting using a probe targeting a region of the *COX1* gene. Under neutral conditions, the mtDNA from all samples migrated as expected for a linear molecule of 16.3 kb, demonstrating that it was full-length and largely free from double-strand DNA (dsDNA) breaks (Fig. 1a). Under denaturing conditions however, the mtDNA separated as a long smear along the length of the lane, indicating that it consisted of single-stranded DNA (ssDNA) fragments ranging from below 250 to over 10,000 nucleotides (nt) in length (Fig. 1b).

**Figure 1.**
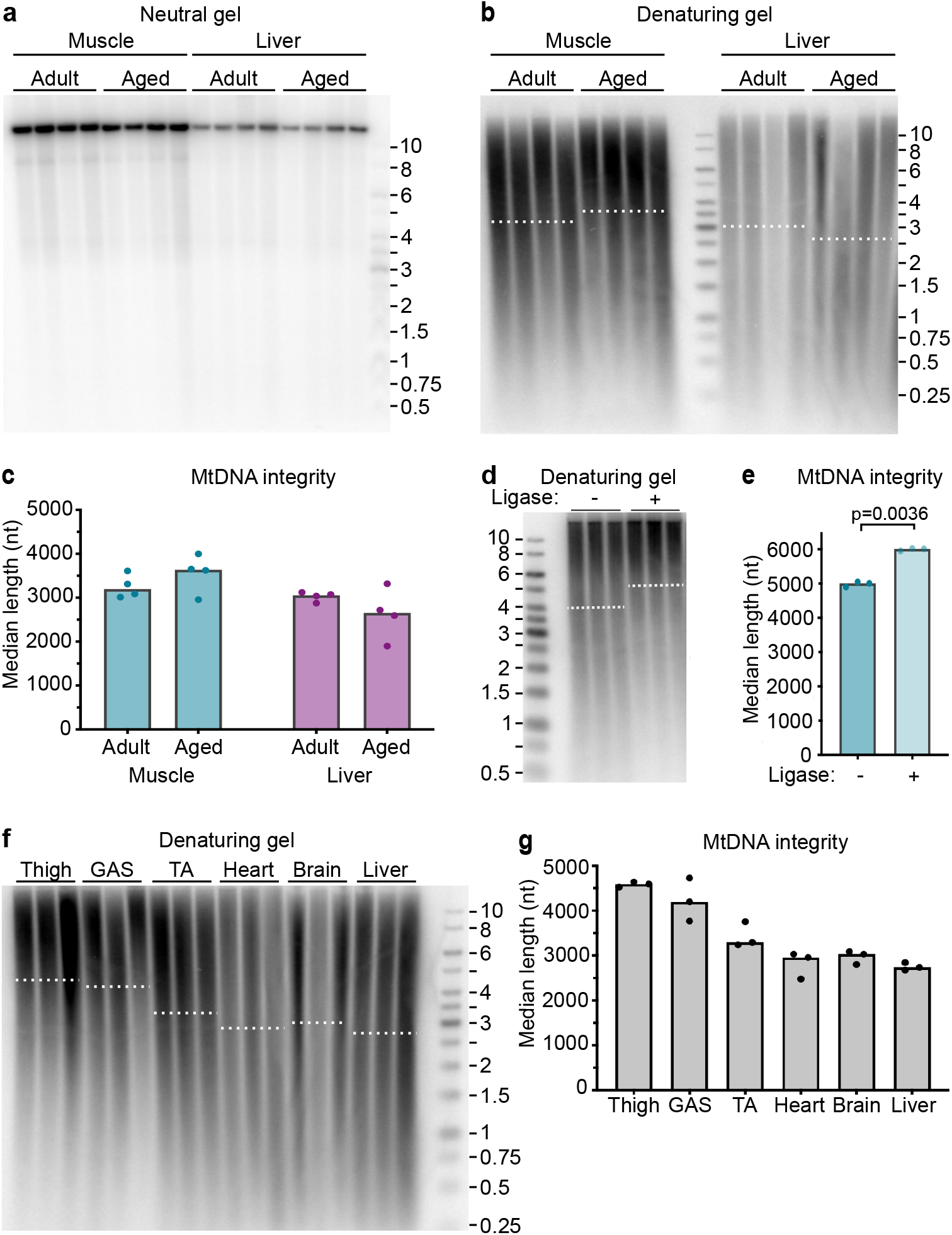
The standard DNA isolation procedure results in mtDNA of low integrity. **a)** DNA isolated from the TA muscle and the liver of 13-16-week-old (*adult*) or 2-year-old (*aged*) animals was linearized with SacI endonuclease and separated on a neutral gel. MtDNA was visualized using a *COX1* probe. Each sample corresponds to an individual mouse. Most mtDNA migrates as expected for a linearized mtDNA molecule; muscle samples also contain some nonlinearized supercoiled molecules that migrate at ca 9 kb. **b)** The same DNA preparations from the TA muscle and the liver of adult or aged animals as in Fig. 1a were separated on a denaturing gel; mtDNA was visualized as above. The signal distribution was quantified and is plotted in Fig. S1a. Dotted white lines represent the median length of mtDNA in each group. **c)** The median length of the mtDNA in Fig. 1b was determined based on the signal distribution (Fig. S1a). The bar indicates the median of the group; each data point represents an individual sample. **d)** Untreated (‘*−*‘) or ligase-treated (‘*+*’) DNA from the TA muscle of adult mice was separated on a denaturing gel and mtDNA visualized by Southern blotting. Dotted lines represent the median of each group. See Fig. S1d for the distribution plot. **e)** The median length of the mtDNA in Fig. 1d was determined based on the signal distribution. The bars indicate the median of a group; each data point represents the median length of an individual sample based on two technical replicates of which a representative one is shown in Fig. 1d. The two groups were compared using a two-tailed paired t-test. **f)** DNA isolated from various tissues of adult mice was separated on a denaturing gel; mtDNA was visualized as above. Each sample corresponds to an individual mouse. The signal distribution was quantified and is plotted in Fig. S1e. Dotted white lines represent the median length of mtDNA in each group. **g)** The median length of the mtDNA shown in Fig. 1f. The bars indicate the median of a group; each data point represents an individual sample. *GAS*, gastrocnemius muscle; *TA*, tibialis anterior muscle. The size of the bands in the DNA ladder are indicated in kb. See also Fig. S1.

To provide a measure of the integrity of the individual strands of mtDNA in the sample preparations, the length distribution of mtDNA in each sample lane of the blot in Fig. 1b was determined relative to size markers run in parallel (Fig. S1a) and expressed as the apparent median length (Fig. 1c). The apparent median length (hereafter referred to as median length) of muscle and liver mtDNA was ca 3,200 and 3,000 nt, respectively, with little variation between the adult and aged groups (Fig. 1c). Notably, the median length of all samples was well below the ca 16,300 nt expected for an intact, full-length strand of mouse mtDNA. Given that the same preparations migrated primarily as full-length mtDNA under neutral conditions (Fig. 1a), the low median length under denaturing conditions was ascribed to the presence of ssDNA breaks that are most likely introduced during sample preparation.

Single-strand DNA breaks can be inflicted by numerous factors, of which DNA shearing, oxidative damage and various endonucleases appear most relevant to our isolation procedure. Shearing and oxidative damage to DNA often generate ‘dirty’ DNA ends that require further processing before ligation can occur (Bertoncini and Meneghini, 1995; Caldecott, 2008; Ohtsubo *et al.*, 2019), whereas nucleases can leave ‘clean’ DNA ends that are directly ligatable. To determine whether the ssDNA breaks observed in our samples contained ligatable ends, we treated the DNA preparations from adult skeletal muscle with T4 DNA ligase and separated ligase-treated and untreated samples on a denaturing gel (Fig. 1d). Ligase treatment resulted in an increase in the median length of the samples (Fig. 1e; distribution plot in Fig. S1d), indicating that the observed low median length was partly due to directly-ligatable ssDNA breaks.

To determine whether the degree of ssDNA breaks was similar across preparations from different tissues, we isolated DNA from a range of mouse tissues using the same standard procedure and examined it on a denaturing gel. Samples from three individual animals were analyzed for each tissue. The integrity of mtDNA in the preparations was found to vary across the tissues analyzed, with highest integrity in skeletal muscle (especially in thigh and gastrocnemius [GAS]), and lower integrity in heart, brain and liver (Fig. 1f-g; distribution plot in Fig. S1e). Considering that all samples were isolated using the same protocol with no differences in handling, and that the ssDNA breaks contained ligatable ends (Fig. 1d-e), we postulated that the variation in the frequency of ssDNA breaks between tissues was likely caused by the differential action of cellular endonucleases. In line with this notion, re-probing of the blot with a probe recognizing nuclear DNA (nDNA) produced a comparable pattern with highest integrity in thigh and GAS, and lowest in liver (Fig. S1f-g), as would be expected if the lower integrity was due to higher activity of endogenous nucleases. Others have previously reported higher endonuclease activity in liver compared to skeletal muscle (Barry et al., 1999). Re-probing of the blots in Fig. 1a-b similarly confirmed that the integrity of nDNA fragments correlated with that of mtDNA (Fig. S1b-c).

Taken together, these findings demonstrate that mtDNA molecules in total DNA preparations prepared by a standard isolation procedure contain several ssDNA breaks per strand of mtDNA, most likely introduced by endogenous nuclease activity during the isolation procedure. These ssDNA breaks are likely to go unnoticed if the quality of DNA preparations is judged only based on neutral gel analysis.

### The frequency of ssDNA breaks can be limited by optimization of the procedure

Strand breaks in the isolated mtDNA can be problematic for downstream analyses requiring high-quality starting material, such as long-range PCR. We therefore sought to improve the integrity of the mtDNA isolated using the standard isolation procedure that involves extraction and freezing of mouse tissues in liquid N_2_, storage at −80°C, overnight lysis of cut tissue at 55°C in buffer containing SDS and proteinase K, phenol-chloroform extraction and ethanol precipitation (see Materials and Methods for details). We reasoned that the highest risk for nuclease action occurred prior to tissue freezing, as well as during the overnight lysis step. Trials were carried out to optimize these two steps using skeletal muscle and liver that contain low and high endogenous nuclease activity, respectively (Fig. 1f-g; (Barry et al., 1999)).

We first analyzed the effect of prompt tissue freezing directly following collection. To this end, the liver and mixed thigh muscles of the hind leg were collected from two euthanized adult animals, divided into five parts of approximately equal weight, and either frozen immediately in liquid N_2_ or incubated at room temperature for 15, 30, 60 or 180 minutes prior to freezing in liquid N_2_. DNA was isolated using the standard procedure and analyzed on a denaturing gel (Fig. 2a; distribution plots in Fig. S2a-b). The median length of mtDNA from the thigh muscles was not obviously affected by the delayed freezing of the tissue (Fig. 2b). In contrast, liver mtDNA exhibited progressively decreasing median length upon delayed freezing of the tissue – while the median length of mtDNA from the promptly-frozen liver was ca 3400 nt, it decreased to 2300 nt following the 180-minute room-temperature incubation (Fig. 2b). As noted previously, the changes in mtDNA integrity were not readily apparent upon neutral gel analysis (Fig. S2c), and the integrity of nDNA fragments followed the same pattern as mtDNA (Fig. S2d). These findings indicate that timely freezing of the tissue can help limit the introduction of ssDNA breaks, particularly in a tissue with high nuclease activity such as the liver.

**Figure 2.**
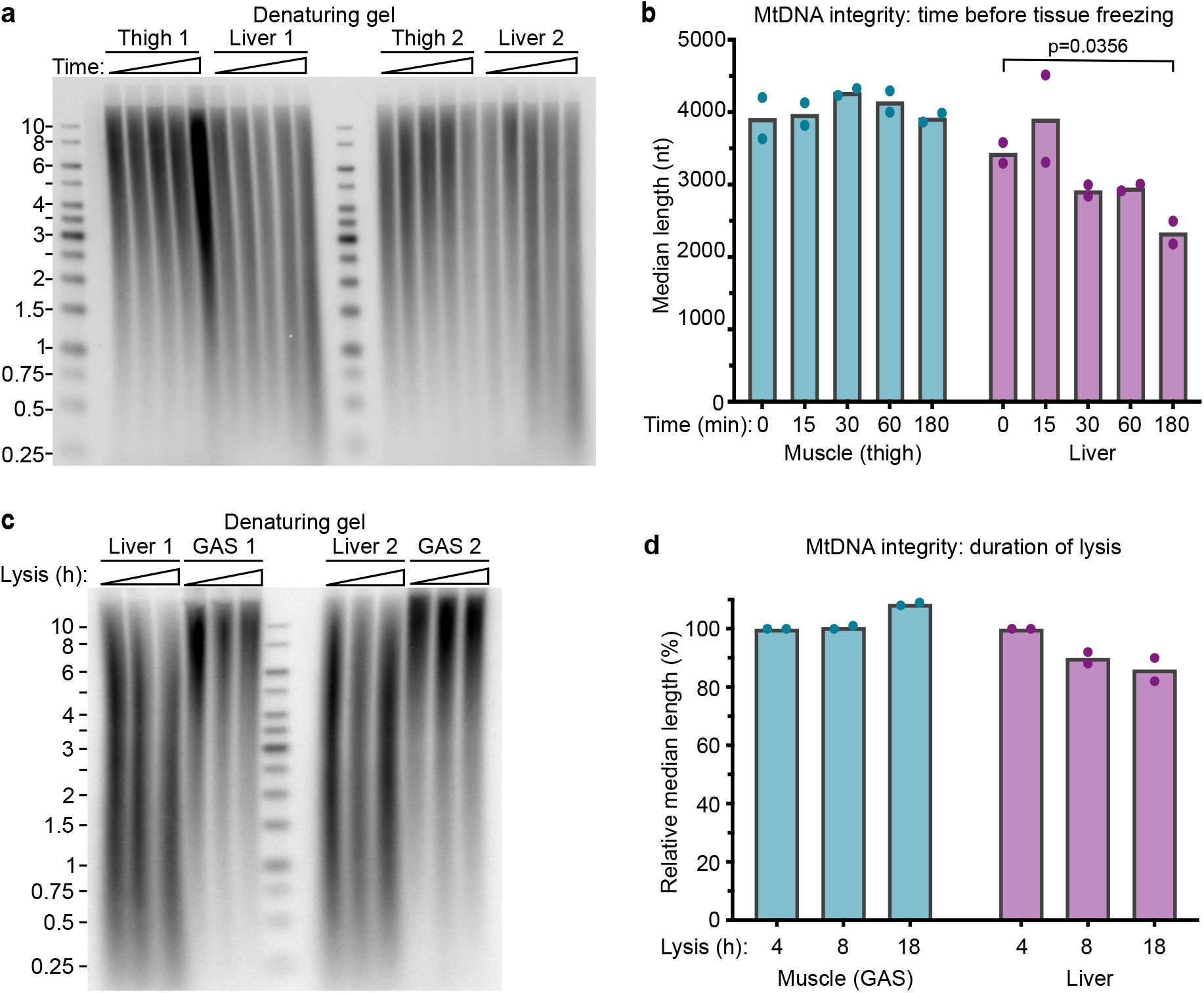
Optimization of the standard isolation procedure can improve the integrity of liver mtDNA. **a)** The thigh muscles and the liver of adult mice were either frozen immediately upon collection, or incubated for 15, 30, 60 or 180 min at room temperature prior to freezing in liquid N_2_. DNA was isolated using the standard procedure, separated on a denaturing gel, and mtDNA was visualized by Southern blotting. The signal distribution was quantified and is plotted in Fig. S2a-b. **b)** The median length of the mtDNA shown in Fig. 2a. The bars indicate the median of a group; each data point represents an individual sample. The 0-min and 180-min timepoints were compared using Welch’s unequal variances t-test; the p-value for the statistically significant comparison is indicated. **c)** Liver and gastrocnemius muscle (*GAS*) was separated into three parts and lysed for 4, 8 or 18 h. DNA was isolated using the standard procedure, separated on a denaturing gel, and mtDNA was visualized by Southern blotting. The signal distribution is plotted in Fig. S2e. **d)** The median length of the mtDNA shown in Fig. 2c is presented relative to that of the 4-h timepoint. The bars indicate the median of a group; each data point represents an individual sample. Variations from the 4-h timepoint are not statistically significant (Welch’s unequal variances t-test). The size of bands in the DNA ladder are indicated in kb. See also Fig. S2.

Next, we considered the possibility that ssDNA breaks are introduced during the overnight lysis of the tissue at 55°C. To test whether limiting the duration of the lysis step could benefit DNA integrity, skeletal muscle (GAS) and the liver from two adult animals were promptly frozen in liquid N_2_ and stored at −80°C. The frozen tissue was divided into three parts of roughly equal weight, and each part was incubated at 55°C in lysis buffer for either 4 h, 8 h or overnight (18 h). DNA was isolated from all samples following the subsequent steps of the standard procedure, and analyzed on a denaturing gel (Fig. 2c; Fig. S2e-f). As observed for delayed tissue freezing, the median length of skeletal muscle mtDNA was not negatively affected by longer lysis (Fig. 2d); in fact, we observed a tendency towards somewhat improved mtDNA integrity after overnight lysis, which may be attributable to improved release of the DNA from this tough tissue following longer lysis. In contrast, the median length of liver mtDNA was highest after only 4 h of lysis, although the differences did not reach statistical significance. Shorter lysis times may therefore benefit the integrity of liver mtDNA. However, 4 h of lysis was insufficient to reach full disruption of the 100 mg of liver used for the extraction, indicating that the optimal duration of the lysis step lies between 4-8 h for liver.

The results of Figure 2 indicate that some improvements in mtDNA integrity can be reached by rapid tissue freezing, and to a lesser extent by limiting the duration of the lysis step, when working with tissues with high nuclease activity such as the liver. However, these measures had little impact on the integrity of mtDNA isolated from skeletal muscle, which — despite better integrity compared to liver mtDNA — still contained several ssDNA breaks per mtDNA strand. Therefore, we next compared the integrity of mtDNA isolated by several alternative DNA isolation protocols with the aim to identify a procedure that yields mtDNA of the highest integrity.

### The integrity of mtDNA isolated by four alternative procedures

We selected four commonly-used methods for isolation of either enriched mtDNA or total DNA from murine solid tissues, and evaluated their effect on mtDNA integrity as judged by migration on both neutral and denaturing gels. Additional variables assessed included DNA yield, mtDNA enrichment (the ratio of mtDNA to nDNA), the duration of the procedure, as well as amenability to scale-up to accommodate a larger number of simultaneous samples. The chosen methods had to allow the simultaneous handling of at least six independent samples and yield a sufficient amount of mtDNA for Southern blot analysis when starting with at most 300 mg of mouse skeletal muscle.

In the first chosen procedure, mitochondria were enriched from tissue homogenates by differential centrifugation and then lysed in the presence of SDS and proteinase K to allow isolation of relatively pure mtDNA (hereafter referred to as ‘pure mtDNA’; see Materials & Methods for experimental details). The other three methods were for the isolation of total DNA: the standard procedure applied in the experiments in Fig. 1–2, where finely-chopped tissue is lysed by overnight incubation with SDS and proteinase K (Sambrook and Russell, 2001); a variant of the standard method where frozen tissue is pulverized in a mortar prior to a 3-h cell lysis; and isolation using the NucleoSpin Tissue kit from Macherey-Nagel. The methods were tested on liver and skeletal muscle. Each individual tissue sample was divided into four parts and DNA isolated by each of the four methods; a total of three samples (biological replicates) was analyzed per method.

We first compared the DNA yield and the enrichment of mtDNA (the ratio of mtDNA to nDNA) isolated by the four methods (Table 1; Fig. 3a). The standard and pulverization methods resulted in highest (total) DNA yields, while the recovery of pure mtDNA was low, only 7.2 and 120 ng DNA per mg of skeletal muscle and liver, respectively (Table 1). When starting with ca 300 mg of skeletal muscle, the total yield is thereby only 2.2 ug of mtDNA. However, the enrichment of mtDNA was high by this method (Fig. 3a). Compared to the standard procedure, the mtDNA from purified mitochondria was enriched *ca* 280 and 570-fold in muscle and liver samples, respectively (Table 1). In terms of yield and enrichment, the pulverization procedure performed similarly to the standard protocol, while the kit-based procedure resulted in an intermediate yield with slight enrichment of mtDNA over nDNA.

**Table 1.** Comparison of DNA yield and mtDNA enrichment of different isolation methods.

|  |  | Average yield<br>(ng of DNA / mg of tissue) | Relative<br>yield* | Relative mtDNA<br>enrichment* |
| --- | --- | --- | --- | --- |
| **Muscle** | **Pure mtDNA** | 7.2 | 0.003 | 280 |
|  | **Pulverized** | 3740 | 1.3 | 1.00 |
|  | **Kit** | 84 | 0.03 | 1.7 |
|  | **Standard** | 2790 | 1 | 1 |
| **Liver** | **Pure mtDNA** | 120 | 0.01 | 570 |
|  | **Pulverized** | 9200 | 1.1 | 0.67 |
|  | **Kit** | 230 | 0.03 | 4.7 |
|  | **Standard** | 8350 | 1 | 1 |

\*Relative to the standard procedure. Non-normalized values for mtDNA enrichment (mtDNA/nDNA) are presented in Fig. 3a.

**Figure 3.**
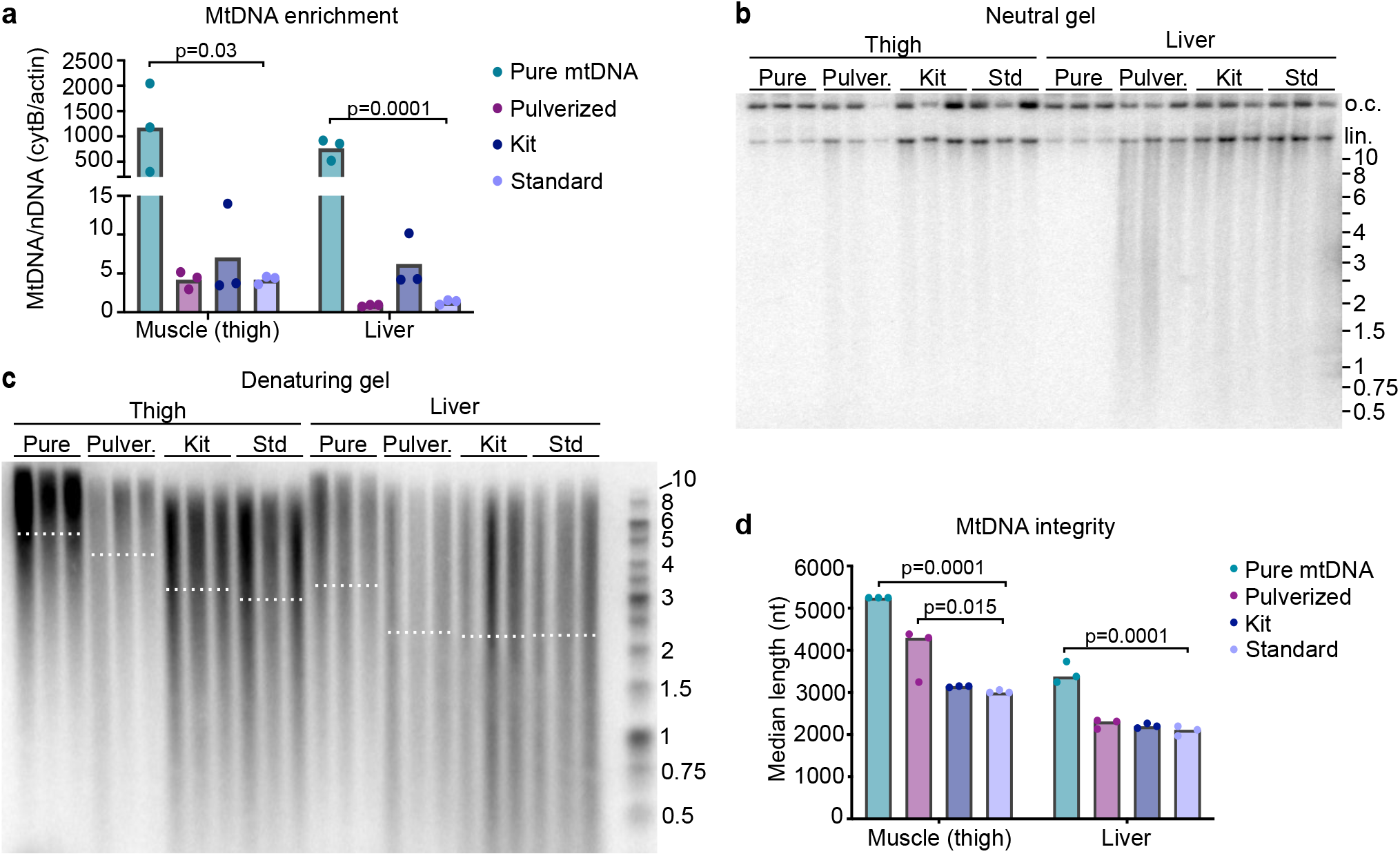
Comparison of mtDNA isolated by different methods. **a)** The enrichment of mtDNA over nDNA in samples prepared from the skeletal muscle (thigh) and liver of adult mice by four different isolation methods was determined by qPCR. The bars indicate the average of each group; each data point represents an individual sample. **b)** The same samples as in Fig. 3a were analyzed on a neutral gel and mtDNA visualized by Southern blotting. Each sample corresponds to an individual mouse. *Pulver.*, pulverized; *Std*, standard procedure; *o.c.*, open circle; *lin*., linear. **c)** The same DNA preparations as in Fig. 3b were separated on a denaturing gel. The signal distribution was quantified and is plotted in Fig. S3a-b. Dotted lines represent the median length of mtDNA in each group. **d)** The median length of the mtDNA from Fig. 3c was determined based on the signal distribution (Fig. S3a-b), and the isolation procedures were compared using one-way ANOVA (n=3). The p-values of statistically significant deviations from the standard procedure are indicated. The bars indicate the median. The size of bands in the DNA ladder are indicated in kb. See also Fig. S3.

Next, the DNA preparations were separated on neutral and denaturing gels, and mtDNA visualized by Southern blotting. Under neutral conditions, some differences in the degree of background smearing could be observed between samples prepared by different procedures; this was most apparent for liver where pure mtDNA samples appeared more intact than the other preparations (Fig. 3b). As the samples were not treated with restriction endonuclease prior to gel analysis, we could also observe some variation in the fraction of mtDNA molecules in open circle *vs*. linear form. The differences in mtDNA integrity between isolation methods were even more pronounced when the same preparations were analyzed under denaturing conditions (Fig. 3c). For both muscle and liver, the pure mtDNA samples migrated noticeably slower than the samples prepared by the standard or kit-based procedures. Accordingly, the median length of mtDNA fragments in the pure mtDNA samples was significantly higher than that of samples prepared by other means (Fig. 3d; distribution plots in Fig. S3a-b). In addition, mtDNA prepared by pulverization of skeletal muscle had a higher median length than that of the kit and standard procedures, while the three total-DNA isolation methods resulted in comparable median lengths of liver mtDNA. We conclude that isolation of DNA from enriched mitochondria preserves the integrity of mtDNA better than the other approaches tested, and is therefore the method of choice when mtDNA of the highest integrity is required.

### The integrity of mtDNA affects the efficiency of downstream applications

As stated before, DNA strand breaks in the extracted mtDNA are expected to decrease the efficiency of sensitive downstream applications such as long-range quantitative PCR where the risk for the DNA polymerase to encounter a strand break in the template strand are substantial. In order to test whether the differences in the integrity of mtDNA isolated by the methods tested in Fig. 3 impact the outcome of long-range quantitative PCR, we amplified a 13.5-kb fragment of mtDNA using the samples as template. A short (145-bp) control fragment was amplified in parallel. As shown in Table 2, the yield of the long 13.5-kb fragment varied between samples isolated by different methods. For both the skeletal muscle and the liver, the 13.5-kb fragment was most efficiently amplified from the pure mtDNA samples that displayed clearly lower C_q_ values than those from the standard isolation method (14.3 and 18.7 for muscle pure mtDNA and the standard method, respectively). In contrast, the C_q_ values for the 145-bp control fragment differed less between methods, as expected for a short amplicon where the chance of encountering a strand break in the template strand is lower (Table 2; 9.3 and 10.0 for muscle pure mtDNA and the standard method, respectively). The improved amplification of the long fragment from pure mtDNA resulted in a smaller difference between the C_q_ values of the long and short fragments (Table 2; right column).

**Table 2.** C_q_ values of a 13.5-kb long fragment and a 145-bp control fragment amplified from mtDNA isolated by various methods.

|  |  | Long fragment C<sub>q</sub> | Short fragment C<sub>q</sub> | C<sub>q</sub><sup>long</sup> - C<sub>q</sub><sup>short</sup> |
| --- | --- | --- | --- | --- |
| **Muscle** | **Pure mtDNA** | 14.3 ± 0.8 | 9.3 ± 0.4 | 4.9 ± 0.9 |
|  | **Pulverized** | 17.5 ± 0.6 | 10.8 ± 0.5 | 6.8 ± 0.8 |
|  | **Kit** | 18.6 ± 0.8 | 11.2 ± 0.6 | 7.4 ± 1.0 |
|  | **Standard** | 18.7 ± 0.4 | 10.0 ± 0.7 | 8.7 ± 0.8 |
| **Liver** | **Pure mtDNA** | 14.5 ± 0.5 | 9.4 ± 0.2 | 5.0 ± 0.5 |
|  | **Pulverized** | 16.9 ± 0.5 | 10.4 ± 0.05 | 6.5 ± 0.5 |
|  | **Kit** | 18.9 ± 0.8 | 10.2 ± 0.1 | 8.7 ± 0.8 |
|  | **Standard** | 18.4 ± 0.6 | 10.6 ± 0.3 | 7.8 ± 0.6 |

Mean ± standard deviation are presented; n=3.

Accordingly, when the quantity of the amplified long fragment was correlated to that of the short fragment, the relative amplification of the long fragment was found to be 16 times higher using the pure mtDNA samples from muscle compared to those isolated by the standard method (Fig. 4). For liver, the pure mtDNA samples showed a 9-fold higher relative amplification than the standard method samples, although the difference did not reach statistical significance. The relative amplification of samples isolated by the pulverization and kit methods did not significantly differ from those of the standard method in either tissue tested, although both methods showed a tendency towards a smaller difference between the C_q_ values of the long and short fragments, suggesting somewhat improved amplification of the long fragment (Fig. 4; Table 2, right column).

**Figure 4.**
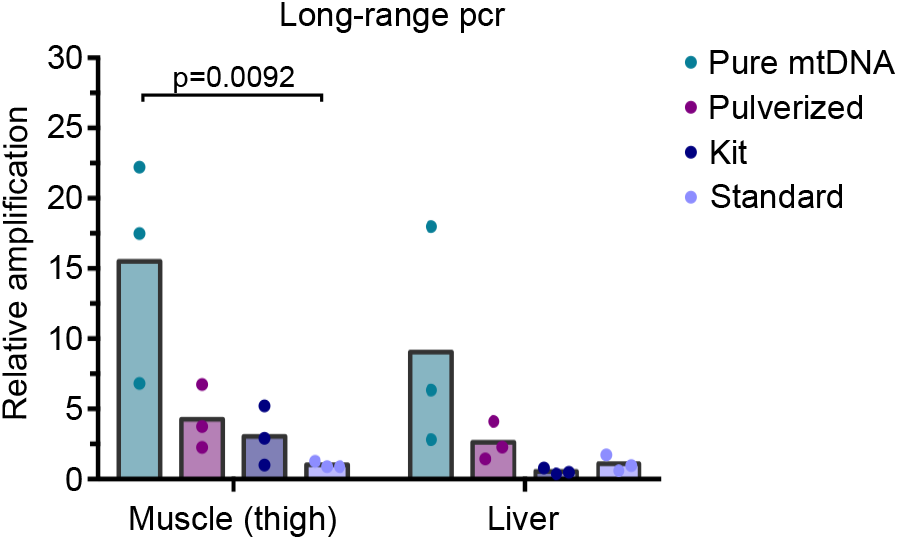
High-integrity mtDNA is a better template for long-range quantitative PCR. The relative amplification of the 13.5-kb amplicon was calculated by normalizing product quantity to that of the 145-bp control amplicon, and the value of the standard method was set to 1. Bars indicate the means of each group. Means were compared to the standard method using one-way ANOVA (n=3). P-values of statistically-significant deviations from the standard method are indicated.

Taken together, our results clearly demonstrate that sample handling and the choice of DNA extraction method influence the integrity of the resulting mtDNA, which in turn can impact the performance of sensitive downstream applications that call for high-integrity starting material. Of the DNA extraction methods tested in this work, the isolation of mtDNA from enriched mitochondria generates a product of highest integrity.

## Conclusions

The integrity of the utilized DNA preparations is a crucial factor for several types of analyses, such as ones based on long-range PCR (Cui et al., 2013; Lehle et al., 2014). The quality of tissue-derived DNA can be jeopardized by the harsh homogenization methods required for disruption of hard tissues such as skeletal muscle, as well as by attack of nucleases prior to their inactivation by the EDTA and proteinase K in the lysis buffer. We find that although the mtDNA prepared by the most commonly-used procedure for isolation of total DNA from mouse tissues appears intact on a neutral gel (Fig. 1a), the median length of the single-stranded mtDNA is well below the expected 16,300 nt (Fig. 1b-c), indicative of the presence of ssDNA breaks. The integrity of mtDNA isolated by the standard procedure varied across tissues, with highest integrity observed from skeletal muscle and lowest from liver, likely due to tissue-specific variations in the activity of endogenous nucleases (Fig. 1f-g). Although the extent of nuclease digestion could be limited somewhat by optimization of the standard DNA isolation procedure (Fig. 2), these measures were insufficient to further improve the integrity of muscle mtDNA. These findings prompted us to search for an alternative mtDNA isolation procedure that better preserves the integrity of mtDNA.

While others have optimized mtDNA extraction for high enrichment of mtDNA over nDNA (Kauppila et al., 2018), we here present a comparison of methods for the isolation of high-integrity mtDNA from two mammalian tissues, skeletal muscle and liver. These two tissues pose different types of challenges for the extraction of high-quality DNA. Skeletal muscle is hard and tough, requiring longer incubation with proteinases to disrupt the tissue, and thus potentially providing a longer time window for nucleases to act before their inactivation. If methods involving mechanical disruption are used, the rougher homogenization required to disrupt muscle cells may bring about DNA shearing, which is a problem especially for the high-molecular-weight nuclear genome. Liver, on the other hand, is soft and thus easy to homogenize, but also one of the tissues with the highest endonuclease activity (Barry et al., 1999). We systematically observed lower integrity of mtDNA from liver than from skeletal muscle, suggesting that the main threat to mtDNA integrity in all our preparations may be cellular nucleases. This was true even for the procedures involving mechanical homogenization (the pure mtDNA and pulverization methods), showing that shearing caused by mechanical homogenization posed a limited risk to mtDNA quality in our experiments.

Of the four methods tested, the ‘pure mtDNA’ method involving extraction of mtDNA from enriched mitochondria yielded mtDNA of the highest integrity from both tissues analyzed. Even this mtDNA is not entirely free of ssDNA nicks, as indicated by a median length below 16,300 nt and by the absence of supercoiled mtDNA forms on the neutral gel (Fig. 3b-d). However, the majority of the pure mtDNA was in open circle conformation, and only a minority contained frequent enough ssDNA breaks to yield a linear mtDNA molecule (Fig. 3b). Accordingly, mtDNA purified by this method showed the highest relative amplification in long-range PCR, making it better-suited for this application than mtDNA isolated by the other methods analyzed in this work. Although it results in the highest mtDNA integrity and enrichment, the pure mtDNA procedure has its disadvantages: it has the lowest yield per mg of tissue used, is the most labor-intensive and difficult to scale up for the parallel handling of many samples (Table 3). In practice, the number of samples simultaneously extracted by this method is limited to ca 8 by the capacity of a suitable high-speed centrifuge and by the time it takes to manually homogenize and handle the samples.

**Table 3.** Summary of the methods assessed for DNA isolation from skeletal muscle.

|  | MtDNA<br>integrity | MtDNA<br>enrichment | Yield | Hands-on time;<br>overall duration* | Suitability for<br>scale-up |
| --- | --- | --- | --- | --- | --- |
| **Pure mtDNA** | High | High | Low | 7 h; 2 d | Low |
| **Pulverized** | Medium | None | High | 4 h; 2 d | Medium |
| **Kit** | Low | Low or none | Medium | 1 h; 1 d | High |
| **Standard** | Low | None | High | 3 h; 3 d | Medium to high |

\*The estimate of overall duration assumes that the re-dissolving of purified DNA is done overnight for all except the kit-based method, as deemed best for optimal solubility.

Of the total DNA isolation procedures, the pulverization method resulted in mtDNA of higher integrity than the standard or kit-based methods when skeletal muscle was used, while for liver mtDNA, a similar integrity was observed with all total DNA isolation methods (Fig. 3c-d). The yield of the pulverization protocol is high, and the method is more amenable to scale-up than the one involving mitochondrial enrichment, although the actual grinding of the frozen tissue can be tedious if numerous samples are to be prepared the same day. Apart from the integrity of muscle mtDNA, the performance of the pulverization method was comparable to the standard procedure that only appreciably surpasses the pulverization method in terms of scalability (Table 3). Finally, the kit-based approach resulted in mtDNA of similar quality as the standard procedure, but with a lower yield. The main advantage of the kit is therefore not the amount or quality of the product but rather the speed and the high suitability of the procedure for scale-up. The number of samples prepared in parallel is limited only by the capacity of the centrifuge so 24 samples can easily be handled simultaneously.

Our results demonstrated that the higher quality of the extracted mtDNA correlates with the amount of product in long-range PCR. In such assays where the readout of the experiment depends on the quality of the mtDNA, the comparison of samples extracted by different isolation methods may therefore introduce unwanted experimental bias and should be avoided. In line with the principles of the 3R’s (replacement, reduction, refinement) in animal experimentation, this work can help reduce the number of experimental animals required for a study by supporting the informed choice of an appropriate DNA isolation procedure from the start of a study (thus limiting the need for repeated analyses), and by decreasing the need for experimental optimization when high-integrity mtDNA is required. Future efforts to further optimize DNA isolation procedures should be aimed at producing mtDNA with fewer nicks even from tissues with high nuclease activity such as the liver.

## Acknowledgements

We gratefully acknowledge Prof. Andrei Chabes for the mouse tissues used in this study. This work was supported by grants from the Swedish Research Council (grant number 2019-01874), The Swedish Cancer Society (grant numbers 19 0022 JIA; 190098 Pj 01 H), The Swedish Society for Medical Research (grant number S17-0023) and the Åke Wiberg Foundation (grant number M20-0132) to P.H.W. The funding sources were not involved in study design, data collection or analysis, or in the decision to publish.

## Author contributions

Experimental design: P.H.W., B.M.R., P.T.; Data collection and analysis: B.M.R., C.M.G., P.T., A.K.N.; Initial draft: P.H.W., B.M.R.; Critical Review: B.M.R., C.M.G., P.T., A.K.N. All authors read and approved the final manuscript.

