## Supplemental data for "The integrity and assay performance of tissue mitochondrial DNA is considerably affected by choice of isolation method"

**Supplemental figure legends**

**Figure S1. Related to Figure 1. a)** The distribution of the signal in each individual lane of Fig. 1b was plotted and used to calculate the apparent median length of the mtDNA (Fig. 1c). The average distribution of biological replicates (n=3) is presented. The size of the DNA on the *x* axis is based on the migration of the DNA ladder run in parallel. **b)** The Southern blot shown in Fig. 1a was stripped and re-probed with a dsDNA probe against the 18S region of nuclear DNA (nDNA). **c)** The Southern blot shown in Fig. 1b was stripped and re-probed with a dsDNA probe against the 18S region of nDNA. **d)** The distribution of the signal in each individual lane of Fig. 1d was quantified and the average distribution of biological replicates (n=3) is presented. The size of the DNA on the *x* axis is based on the migration of the DNA ladder run in parallel. **e)** The distribution of the signal in each individual lane of Fig. 1f was quantified and the average distribution of biological replicates (n=3) is presented. The size of the DNA on the *x* axis is based on the migration of the DNA ladder run in parallel. *GAS*, gastrocnemius muscle; *TA*, tibialis anterior muscle. **f)** The Southern blot shown in Fig. 1f was stripped and re-probed with a dsDNA probe against the 18S region of nDNA. **g)** The distribution of the signal in each individual lane of Fig. S1f was plotted and used to estimate the integrity of the nDNA fragments that had run into the gel. The average signal distribution of biological replicates (n=3) is presented. The size of the DNA on the *x* axis is based on the migration of the DNA ladder run in parallel.

**Figure S2. Related to Figure 2. a)** The distribution of the signal in each individual lane with thigh sample in Fig. 2a was plotted and used to calculate the apparent median length of the mtDNA (Fig. 2b). The average distribution of biological replicates (n=2) is presented. **b)** The distribution of the signal in each individual lane with liver sample in Fig. 2a was plotted and used to calculate the apparent median length of the mtDNA (Fig. 2b). The average distribution of biological replicates (n=2) is presented. **c)** The DNA shown in Fig. 2a was linearized by digestion with SacI and separated on a neutral gel; mtDNA was visualized using a dsDNA probe for the *COX1* region. *Uncut*, a sample not treated with SacI; *o.c.*, open circle; *lin.*, linear. **d)** The Southern blot shown in Fig. 2a was stripped and re-probed with a dsDNA probe against the 18S region of nuclear DNA (nDNA). **e)** The distribution of the signal in each individual lane of Fig. 2c was quantified and the average distribution of biological replicates (n=2) is presented. **f)** The Southern blot shown in Fig. 2c was stripped and re-probed with a dsDNA probe against the 18S region of nuclear DNA (nDNA).

**Figure S3. Related to Figure 3. a)** The distribution of the signal in each individual skeletal muscle-containing lane from Fig. 3c was plotted and used to calculate the apparent median length of the mtDNA (Fig. 3d). The average distribution of biological replicates (n=3) is presented. The size of the DNA on the *x* axis is based on the migration of the DNA ladder run in parallel. **b)** The distribution of the signal in each individual lane of liver mtDNA from Fig. 3c was plotted and is presented as in Fig. S3a.

**Supplemental Figure 1; related to Figure 1.**

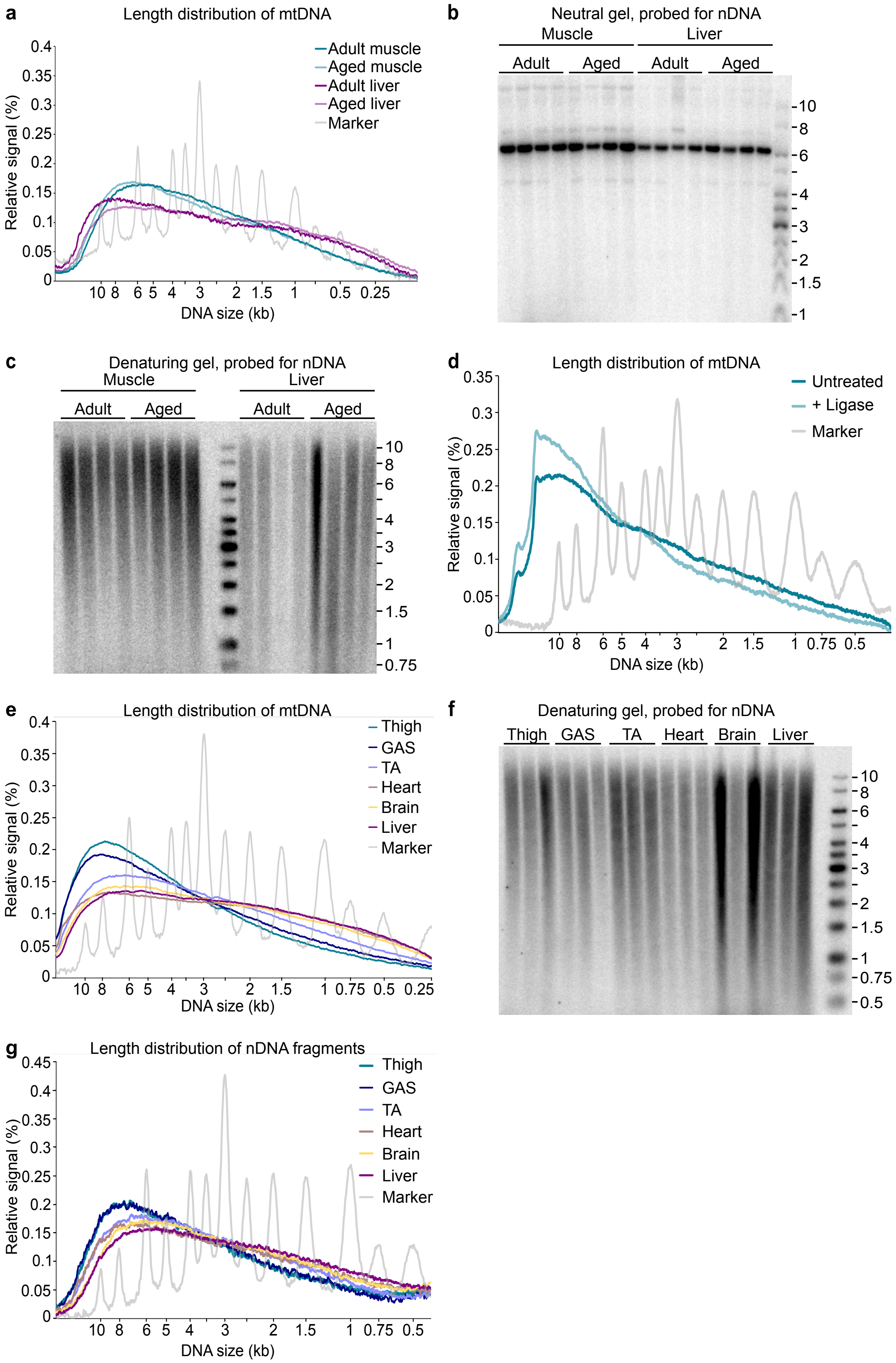

**Supplemental Figure 2; related to Figure 2.**

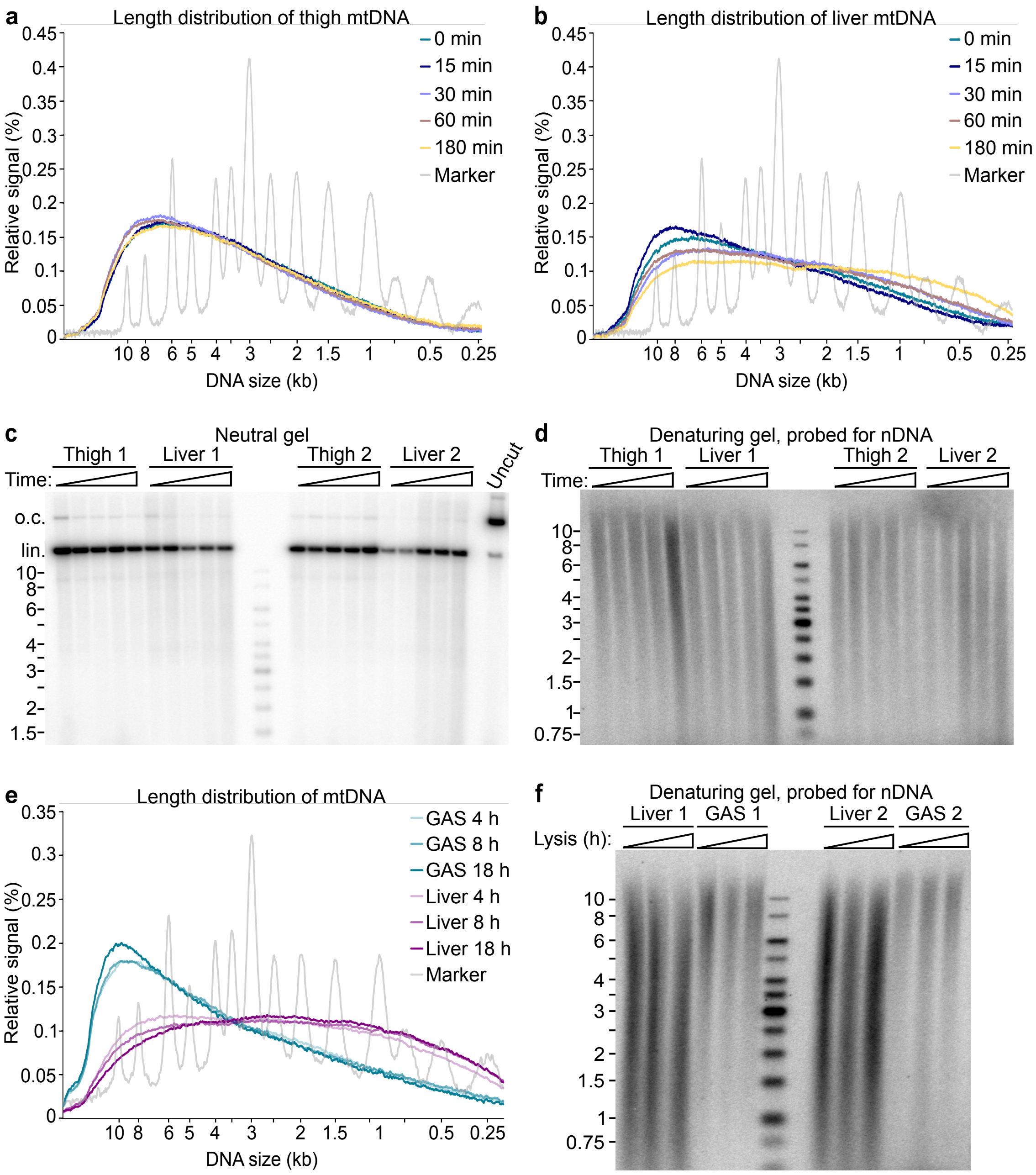

**Supplemental Figure 3; related to Figure 3.**

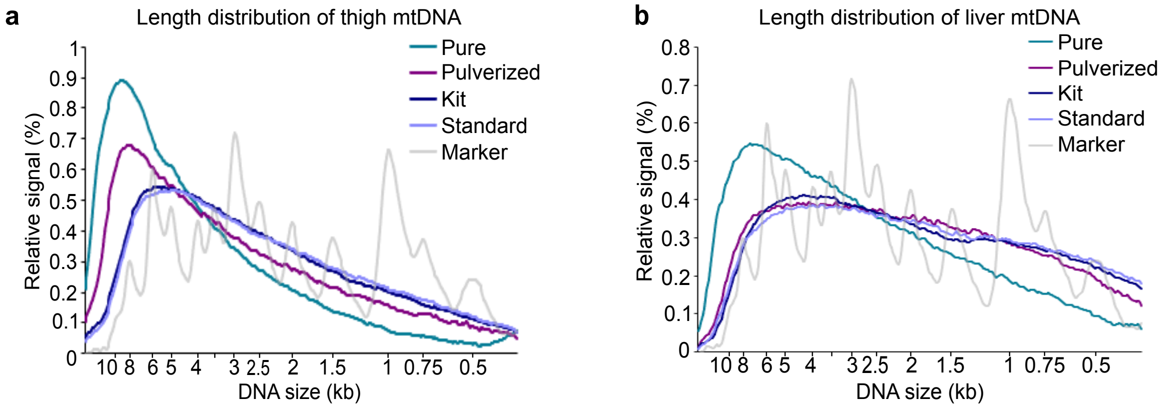
